# Analysis of the effects of related fingerprints on molecular similarity using an eigenvalue entropy approach

**DOI:** 10.1101/853762

**Authors:** Hiroyuki Kuwahara, Xin Gao

## Abstract

Two-dimensional (2D) chemical fingerprints are widely used as numerical features for the quantification of structural similarity of chemical compounds, which is an important step in similarity-based virtual screening (VS). Here, using an eigenvalue-based entropy approach, we sought to identify 2D fingerprints with little to no contribution to shaping the eigenvalue distribution of the feature matrix as related fingerprints and examined the degree to which these related 2D fingerprints influence molecular similarity scores via Tanimoto coefficient. We found that there are many related fingerprints in publicly available fingerprint schemes and that their presence in the feature set tends to decrease the similarity scores. Our results have implication in the optimal selection of 2D fingerprints and the identification of potential hits for compounds with target biological activity in VS.

## 1 Introduction

Virtual screening (VS) is a computational approach that is widely used as a cost-effective alternative to the traditional high-throughput screening for the selection of initial hits in a search for drugs with a given biological activity [9,13]. The foundation of similarity-based VS is structure-activity relationship (SAR), a concept in which molecules with similar structures are destined to have similar biological activities. In such VS applications, thus, the quantification of structural similarity of molecules is a crucial step. To quantify the structural similarity of a pair of molecules, the Tanimoto similarity measure is commonly applied to fingerprint features based on their two-dimensional (2D) structures. These 2D fingerprints represent each molecule as a binary (0 or 1) vector characterizing the absence or the presence of specific properties of its 2D structure. Although this feature representation is simple, it has been reported to be more effective than those using more complex features such as 3D structural patterns [12, 15].

There are libraries of predefined 2D chemical fingerprint dictionaries available to represent molecules as binary vectors [3]. Among the most commonly used fingerprint schemes for similarity quantification is molecular access system (MACCS) [4], which was reported to cover many useful 2D features for virtual screening [10]. While these predefined fingerprint dictionaries are easy to use, previous studies demonstrated that the selection of relevant 2D fingerprints from the original set resulted in better performance [2,6,7]. These feature selection methods typically focus on supervised machine learning settings in which to select a subset of relevant 2D fingerprints that intend to enhance the generality to discriminate chemical compounds with a given biological activity against those without. For example, Nisius, et al. ranked 2D fingerprints by applying the Kullback-Leibler divergence to each fingerprint to quantify its asymmetric usage between the active compound class and the inactive one [11]. Given the nature of drug discovery, however, these supervised feature selection approaches inevitably face a challenging class imbalance problem in practice as available compounds with the target biological activity is most likely very scarce. That is, had the number of target bioactive compounds been large enough to begin with, a pipeline to discover more of the same would not have probably warranted a large cost of investment.

Here, we focus on a different issue in the combination of 2D fingerprints and analyze the effects of related fingerprints on the quantification of molecular similarity using eigenvalue-based entropy. The eigenvalue-based entropy was introduced by Alter et al. [1] to indicate the weight distribution of gene expression eigenvectors for analysis of temporal gene expression patterns. Varshavsky, et al. [14] developed an unsupervised feature selection method which ranks each feature by measuring its contribution to the eigenvalue-based entropy. We define the relatedness of each 2D fingerprint based on the degree to which the shape of the eigenvalue distribution of the feature matrix is changed. And, by using the eigenvalue-based entropy as the scaler value to indicate the distribution of eigenvalues, we determined related 2D fingerprints. Thus, we define a related 2D fingerprint as a feature which has a (quasi) linear relationship with some other fingerprints in the feature set regardless of its relevance and importance for the discriminability. In this paper, we applied MACCS and Pubchem fingerprint schemes to a human metabolite dataset and identified up to 63% of the total fingerprints as related ones. We found that the presence of related 2D fingerprints has a tendency to lower similarity scores. Our analysis demonstrated that these effects can result in a decrease in the number of potential hits and qualitatively change the outcome of VS.

## 2 Methods

### 2.1 Datasets

From Human Metabolome Database (HMDB) [17], we retrieved 2D structure data for 25,376 metabolites on September 5, 2019. With a filtering for the metabolites found in blood with the metabolite status being “Detected and Quantified,” we further obtained the information about 3202 metabolites for the blood specimen.

### 2.2 Molecular similarity measure

We used the implementation of CDK (version 2.3) [16] to compute 166-bit MACCS and 881-bit Pubchem fingerprint vectors. To measure the similarity of a pair of compounds, *a* and *b*, we computed Tanimoto coefficient of their *l*-bit fingerprint vectors, *v*_*a*_ and *v*_*b*_ as follows:

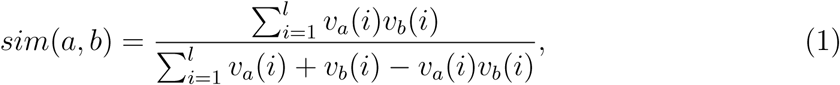

where *v*(*i*) represents the *i*-th element of vector *v*.

### 2.3 Eigenvalue-based entropy

Let *A* be an *m* by *n* matrix. Then, an *n* by *n* symmetric matrix *A*^*T*^ *A* is positive semidefinite and has real eigenvalues *λ*_1_ ≥ *λ*_2_ ≥ … ≥ *λ*_*n*_ ≥ 0. By defining *q*_*j*_ to be the *j*-th normalized eigenvalue 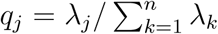, we computed a single value which indicates the complexity of the distribution of eigenvalues with the normalized entropy of eigenvalues [1] as follows:

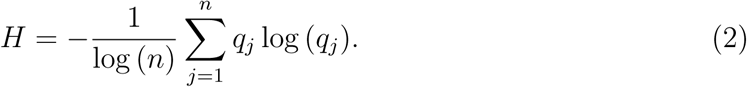

This entropy ranges from 0 to 1, with 0 indicating that the dataset can be constructed based on a single eigenvector and 1 indicating that each eigenvector has an equal contribution to the dataset. Eigenvalues were computed using the *svd* function in R to compute eigenvalues.

### 2.4 Eigenvalue-based fingerprint contribution measure

Suppose we have *m* compounds, each of which is expressed with *n*-bit 2D fingerprints. That is, we have an *m* by *n* matrix *A* whose element *a*_*i,j*_ represents the value of the *j*-th fingerprint for the *i*-th compound. Let *A*[−*i*] be an *m* by *n* matrix that has all but the *i*-th column of *A* with the *i*-th column replaced by a zero column. We computed the contribution of the *i*-th (1 ≤ *i* ≤ *n*) fingerprint, *h*_*i*_ as *h*_*i*_ = *H*(*A*[−*i*]) where *H*(*M*) is the eigenvalue-based entropy of matrix *M* given by Equation 2. Note that, because we can first compute *n*-by-*n* matrix from *A*^*T*^ *A*, the computation of each fingerprint entropy depends on the number of the fingerprints and not on the number of compounds, which is presumed to be very large.

## 3 Results

### 3.1 Presence of highly correlated fingerprints

MACCS keys are 166-bit 2D structure fingerprints that are commonly used for the measure of molecular similarity. Because each bit is either on (i.e., 1) or off (i.e., 0), MACCS 166 keys can represent more than 9.3 × 10^49^ distinct fingerprint vectors. We generated 24,921 MACCS fingerprint vectors using the metabolite data we obtained from HMDB [17] (see Methods). After filtering out duplicates, we ended up with 3,125 unique fingerprint vectors. On average, thus, 8 metabolites were represented by the same MACCS fingerprint vector, indicating a high degree of collisions. The high level of collided metabolites suggests the possibility that many MACCS keys describe related 2D substructure characteristics.

To analyze the use of each MACCS key, we first counted the occurrence of on bit for each key in the 3,125 unique fingerprint vectors (Fig. 1A). We found that 39% of the 166 MACCS keys are on (i.e., 1) for fewer than 10% of the fingerprint vectors, while only 1 key is on for more than 90% of the vectors. This skewed use of molecular fingerprints (*γ*_1_ = 0.777) indicates that many fingerprint bits are set to be off (i.e., 0) in most of the vectors, resulting in highly similar usage patterns. However, since Tanimoto coefficient, the most commonly used 2D fingerprint-based similarity measure, does not consider the off bits (see Methods), its similarity analysis of these HMDB metabolites may not be influenced by the fingerprints with many off bits.

**Figure 1.**
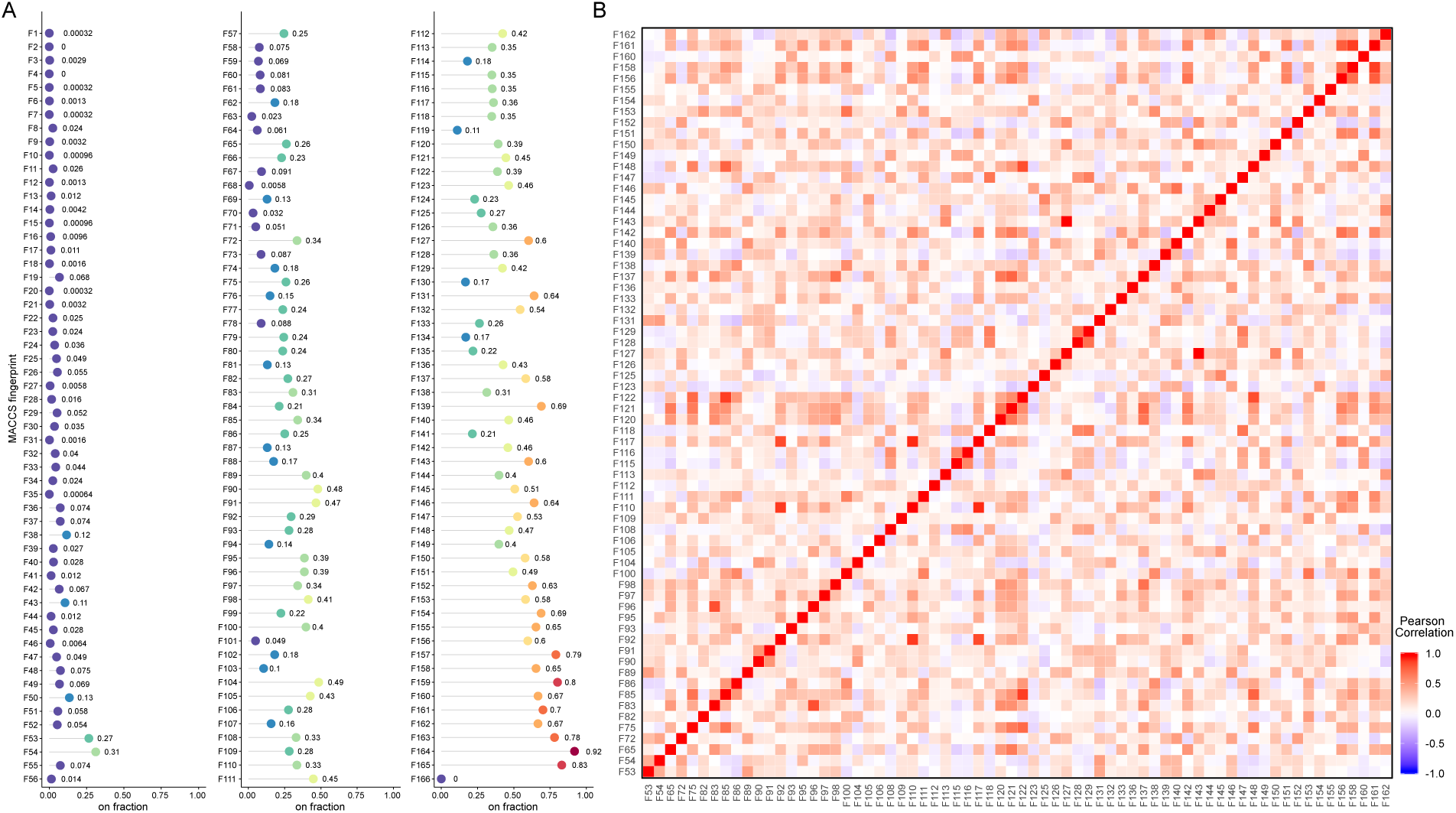
Fingerprint usage patterns of MACCS 166 keys on HMDB metabolite dataset. (A) The on-bit count of each key. (B) The pairwise Pearson’s correlation coefficient value for each pair of 68 MACCS keys with moderate on-bit counts.

We next analyzed the association of MACCS keys whose on-bit counts are more moderate and whose effects on similarity measure are assumed to be more profound. To this end, we focused on a subset of the MACCS keys whose on-bit counts are in a range between 25% and 75% of the total number of the unique vectors. We obtained 68 MACCS keys that satisfied this constraint and computed their pairwise correlation coefficient values (Fig. 1B). We found that a large fraction of the pairs (73%) had positive correlation (*r* ≥ 0). Out of 2,278 fingerprint pairs, while none had strong negative correlation (*r* ≤ −0.5), 105 had strong positive correlation (*r* ≥ 0.5). Among these positively correlated pairs, the 127th and the 143rd fingerprints, both of which had 1,875 on-bit counts, had the perfect positive correlation, suggesting that 2D structures characterized by these two fingerprints are highly related. Although the correlation coefficient can capture only a limited type of related fingerprints, these results suggest the prevalence of related fingerprints in the predefined 2D fingerprint dictionaries.

### 3.2 Characterization of related fingerprints

We next sought to analyze the extent to which more general types of related fingerprints were present in the MACCS and Pubchem fingerprint dictionaries. To this end, we gathered 3202 metabolites found in the blood specimen from the HMDB metabolite dataset and filtered out compounds with duplicate 2D structures, duplicate fingerprint vectors, and all-zero MACCS fingerprint vectors. In addition, we removed each compound whose MACCS fingerprint vector has off bits for more than 90% of the fingerprints.

With this data preprocessing, we selected 1023 metabolites that have unique fingerprint vectors. The 1023 by 166 matrix formed with the MACCS fingerprints had the rank of 144, where the column represents the fingerprints and the row represents the metabolite (see Methods), indicating that the pattern of ∼15% of MACCS fingerprints can be completely captured by the rest. The Pubchem fingerprints resulted in a 1023 by 881 fingerprint matrix which had the rank of 377, indicating even more pronounced effects of rank deficiency with more than half of the fingerprints completely characterized by linear combinations of 377 fingerprints.

To assess the degree of related fingerprints, we defined the relatedness using the eigenvalue-based entropy (see Methods). This eigenvalue-based entropy measure indicates the shape of the eigenvalue distribution [1], with its value ranging from 0 to 1 where a higher value indicates that the matrix can be reconstructed with a linear combination of a smaller number of eigenvectors. The distribution of the normalized eigenvalues for the MACCS and Pubchem fingerprint matrices shows that the first component has a high weight in both (Fig. 2A), indicating that their entropy values be lower. Indeed, the entropy values of the original MACCS and Pubchem matrices were 0.474 and 0.355, respectively. To measure the relatedness of the *i*-th fingerprint with the other fingerprints, we computed the change in the entropy between the original fingerprint-feature matrix and the feature matrix without the *i*-th fingerprint (see Methods). This can indicate the contribution of the *i*-th fingerprint to shaping the eigenvalue distribution, which, in turn, allows us to evaluate the degree to which the *i*-th fingerprint is linearly related to some other fingerprints. Figure 2B shows the distribution of the fingerprint entropy values for both the MACCS and Pubchem schemes. We found that there is a high peak at the entropy value of the original fingerprint-feature matrix with many fingerprints having their entropies in near the original one, indicating that these fingerprints do not contribute much to the eigenvalue distribution and are highly related to some other fingerprints.

**Figure 2.**
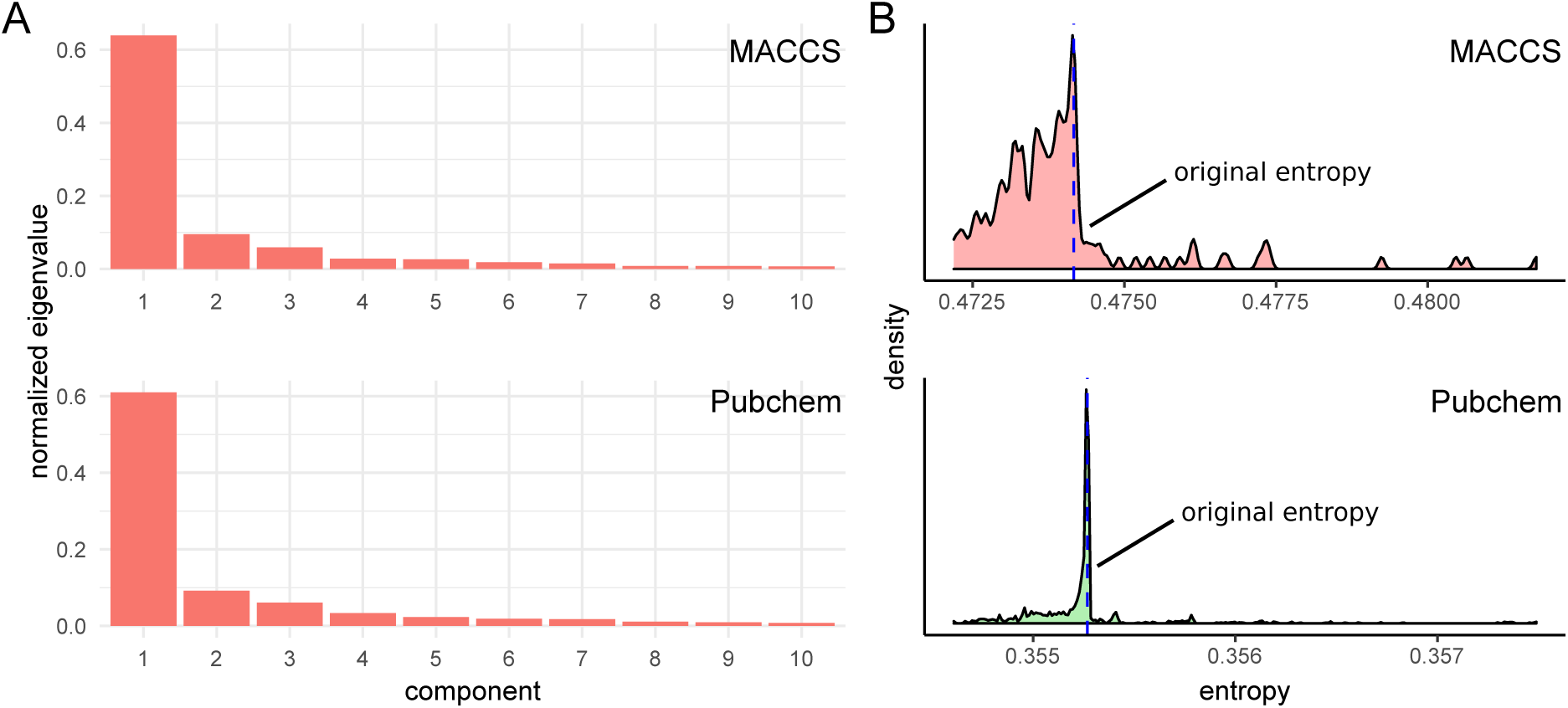
Eigenvalue-based analysis of MACCS and Pubchem fingerprint matrices. (A) Normalized eigenvalues of the first 10 components for MACCS and Pubchem fingerprint matrices. (B) The distribution of the eigenvalue-based entropy for MACCS and Pubchem fingerprints.

By measuring the relatedness based on the distance between the original entropy and the entropy for each fingerprint, we selected related fingerprints from the original MACCS and Pubchem fingerprint dictionaries. Let *h*_0_ and *h*_*i*_ (1 ≤ *i* ≤ *n*) be the eigenvalue-based entropy of the original feature matrix and for the *i*-th fingerprint, respectively. Then, we selected the *i*-th fingerprint as a related feature if *h*_*i*_ satisfies the following condition:

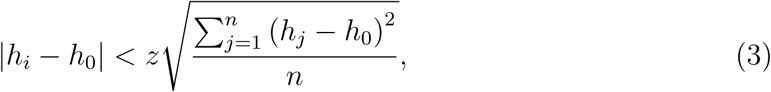

where *z* is a reduced-level threshold parameter, which was set to 0.1, 0.2, and 0.3 in this study. Based on this approach, we found that many fingerprints in the MACCS and Pubchem dictionaries are related (Fig. 3). With the reduced level threshold being 0.1, 0.2, and 0.3, we identified 28, 48, 62 related fingerprints in MACCS and 454, 525, and 555 in Pubchem, respectively. A larger fraction of the related fingerprints identified in the Pubchem scheme with the low threshold value was expected given that the distribution of its fingerprint entropies had a higher density of the fingerprints near the original entropy.

**Figure 3.**
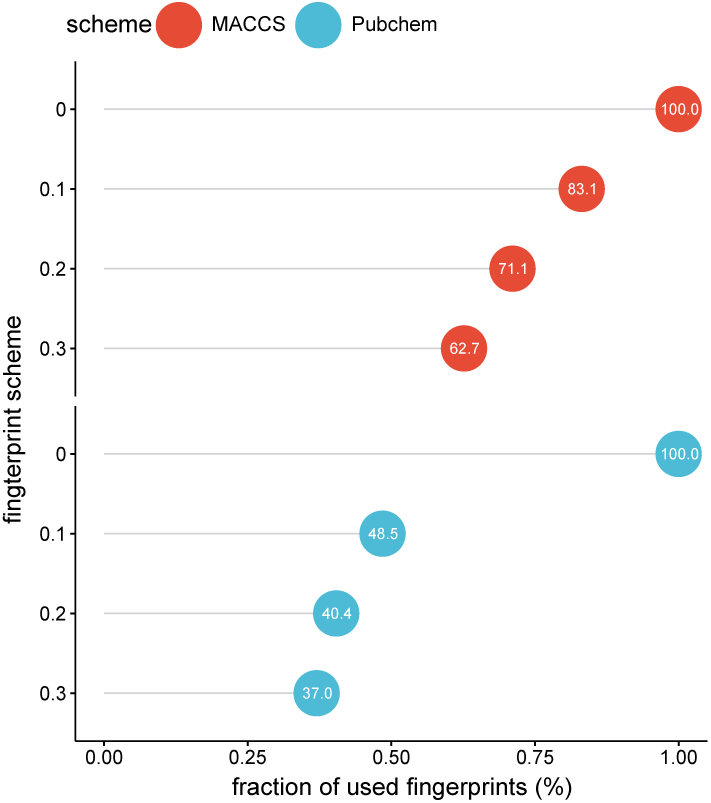
The fraction of used fingerprints with respect to given reduced levels for MACCS and Pubchem fingerprints. The reduced level 0 indicates the fraction for the original fingerprints.

### Effects of related fingerprints to decrease molecular similarity

Given the subsets of fingerprints that we obtained through the use of the fingerprint-based entropy, we analyzed the effects of removing related fingerprints on the Tanimoto similarity score. To this end, we first constructed 1023 by 1023 similarity matrix by computing the similarity score for each metabolite pair and generated the normalized eigenvalues of similarity matrices (Fig. 4A). The comparison of the first six components suggests that the similarity matrices computed from fingerprint sets with various reduction levels in both MACCS and Pubchem schemes are similar. Next, we measured the absolute difference of 522,753 distinct metabolite pairs between the original fingerprint set and reduced fingerprint sets. We found that the difference increases as the reduced level threshold increases in both MACCS and Pubchem fingerprint dictionaries (Fig. 4B). While both fingerprint schemes had quantitatively similar levels of absolute differences with the reduced level at 0.01, the difference became wider as the threshold increases particularly in MACCS, suggesting the effects of removing related fingerprints were greater in the MACCS scheme even though a higher fraction of fingerprints were removed in the Pubchem scheme.

**Figure 4.**
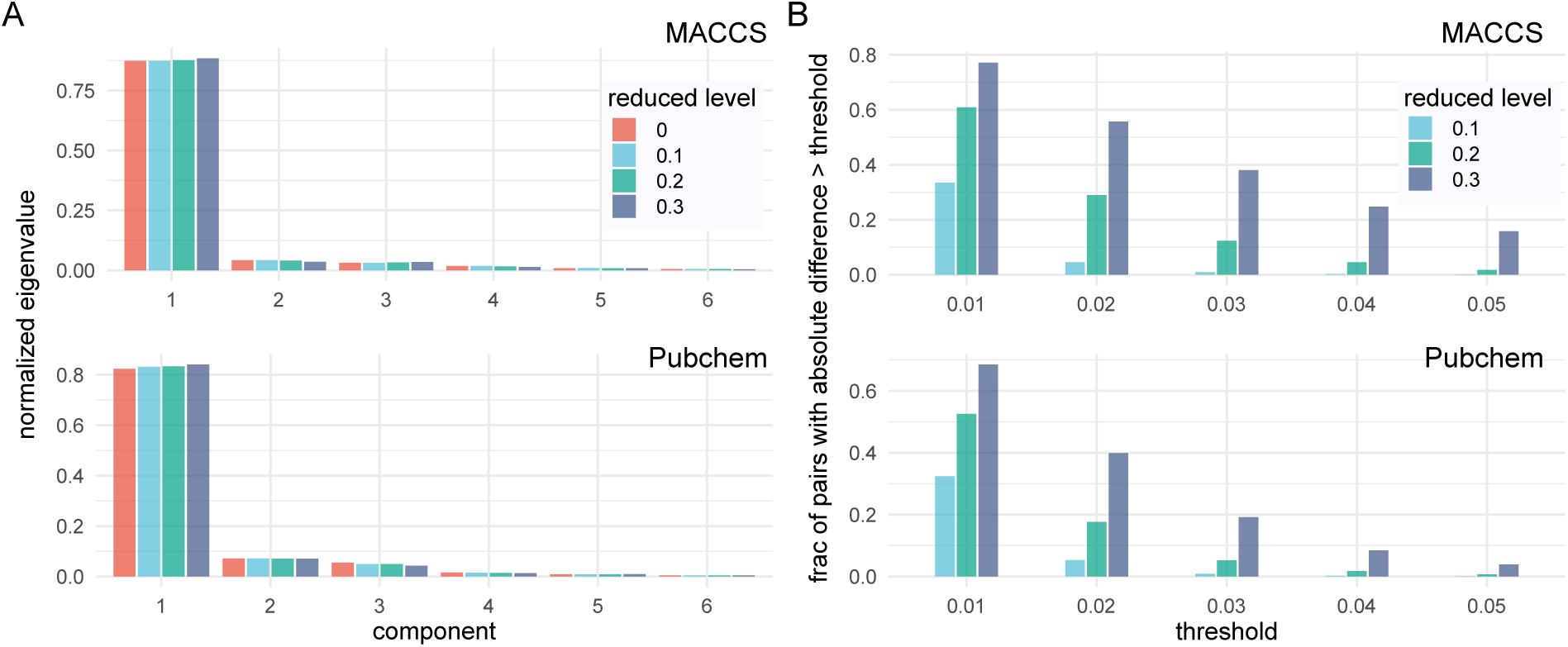
Comparison of the Tanimoto similarity scores based on different reduction levels. (A) The comparison of the first 6 principal components with respect to different reduction levels. (B) The comparison of the fraction of metabolite pairs with the absolute similarity score difference between the original set of fingerprints and a given reduced set of fingerprints exceeding specified threshold values.

**Figure 5.**
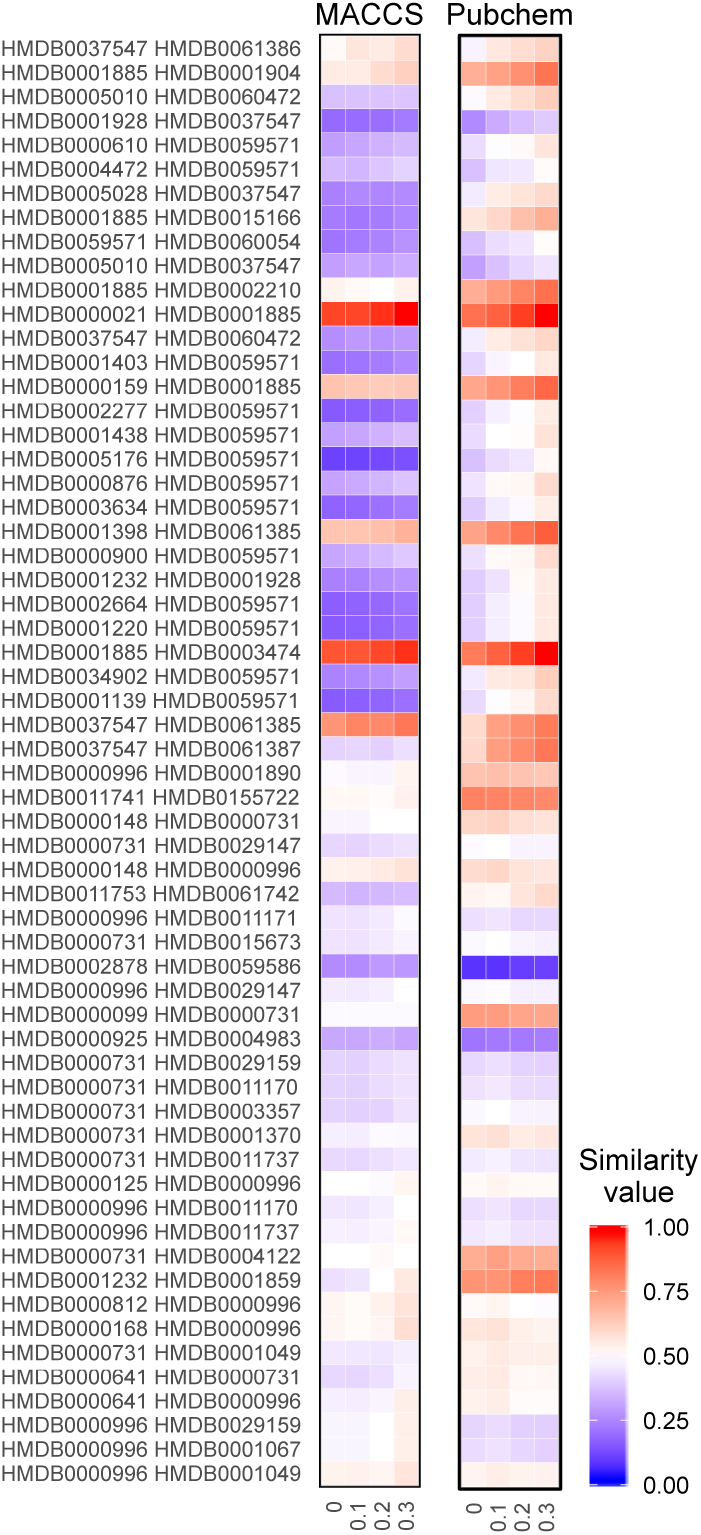
Illustration of 60 metabolite pairs with high levels of changes in Tanimoto similarity measures. Heatmap showing the similarity scores of 60 metabolite pairs based (y-axis) on given levels of reduced fingerprint sets (x-axis). From the MACCS and Pubchem fingerprint dictionaries, 30 pairs are selected from each based on the difference between the original set of fingerprints and a reduced set of fingerprints with reduced level 0.3.

To further analyze the effects of related fingerprints on the Tanimoto similarity scores, we grouped the metabolites in the blood specimen into four classes: drug; microbial; plant; and endogenous, using metabolite information retrieved from HMDB. In each of these four categories, we computed the average of the pairwise Tanimoto similarity scores. The results show that the average similarity scores from the reduced fingerprint sets are quantitatively close to those from the original fingerprint sets (Table 1). We also found that similarity scores from the reduced fingerprint sets tend to be marginally higher than those from the original fingerprint sets. In other words, the inclusion of related fingerprints has negative effects and tends to slightly decrease the Tanimoto similarity score. This trend is stronger in the Pubchem fingerprint scheme in which the fingerprint sets from all of the three reduced levels resulted in higher similarity scores than the original one in all of the four subclasses.

**Table 1.** The average Tanimoto similarity score for five classes of metabolites in the blood specimen for the MACCS and Pubchem fingerprint schemes with different reduction levels.

| scheme | level | drug | microbial | plant | endogenous | all |
| --- | --- | --- | --- | --- | --- | --- |
| MACCS | 0 | 0.3008 | 0.3531 | 0.3662 | 0.3211 | 0.3142 |
| MACCS | 0.1 | 0.2990 | 0.3489 | 0.3696 | 0.3190 | 0.3122 |
| MACCS | 0.2 | 0.2944 | 0.3536 | 0.3723 | 0.3188 | 0.3118 |
| MACCS | 0.3 | 0.3013 | 0.3578 | 0.3785 | 0.3211 | 0.3149 |
| Pubchem | 0 | 0.3048 | 0.3323 | 0.3873 | 0.2967 | 0.2968 |
| Pubchem | 0.1 | 0.3104 | 0.3390 | 0.3922 | 0.3012 | 0.3016 |
| Pubchem | 0.2 | 0.3167 | 0.3384 | 0.3974 | 0.3043 | 0.3053 |
| Pubchem | 0.3 | 0.3217 | 0.3395 | 0.4010 | 0.3071 | 0.3085 |

### Removal of related fingerprints to increase similarity accuracy

Although our analysis showed the effects of related fingerprints on the Tanimoto similarity measures, it is not clear if those effects can lead to qualitatively significant changes in structural similarity-based molecule screening. To analyze potential effects of related fingerprints in such SAR-based analysis, we used a dataset consisting of 100 drug-like molecular pairs from DrugBank 3.0 [8], which 143 experts analyzed to provide their yes or no binary decisions about the structural similarity [5]. To serve this dataset as the correct reference for similar compounds, we selected a subset of the 100 pairs whose similarity was supported by at least 80% of the experts, resulting in 33 pairs of similar compounds.

Using this dataset, we analyzed the performance of each fingerprint set by computing three measures: the mean similarity value; the number of pairs with high similarity values (i.e., *sim* ≥ 0.8); and the number of pairs with low similarity values (i.e., *sim <* 0.5). We decided to only focus on these measures because in virtual screening molecular similarity is used to filter out incompatible compounds and to generate initial hits with potentially similar bioactive properties in order to capture them in follow-up screenings [13]. That is, in the filtering for the initial hits, as long as true positives are included, the number of false positives is not as important.

Table 2 shows the results of our analysis. We found that in both MACCS and Pubchem schemes, the reduced sets of fingerprints increased the average similarity scores of the 33 pairs. The number of high similarity sores also increased by removing related fingerprints; in the MACCS scheme, the original fingerprint set and the reduced fingerprint set with threshold 0.03 resulted in 73% and 76% of the compound pairs with high similarity scores, while in the Pubchem scheme, they resulted in 88% and 94% of the pairs with high similarity scores. Furthermore, while the original fingerprint set and the reduced fingerprint sets of the Pubchem scheme did not result in any compound pairs with low similarity scores, in the MACCS scheme, the original fingerprint set had one pair with a low similarity score, which had high enough scores in all of the reduced fingerprint sets.

**Table 2.** Summary of the results from 33 pairs of similar compounds with a high consensus from 143 experts.

| scheme | level | mean | high sim<sup>a</sup> | low sim<sup>b</sup> |
| --- | --- | --- | --- | --- |
| MACCS | 0.0 | 0.8724 | 24 | 1 |
| MACCS | 0.1 | 0.8745 | 24 | 0 |
| MACCS | 0.2 | 0.8748 | 24 | 0 |
| MACCS | 0.3 | 0.8801 | 25 | 0 |
| Pubchem | 0.0 | 0.9163 | 29 | 0 |
| Pubchem | 0.1 | 0.9224 | 30 | 0 |
| Pubchem | 0.2 | 0.9282 | 30 | 0 |
| Pubchem | 0.3 | 0.9334 | 31 | 0 |

<sup>a</sup>The number of instances in which the similarity score is greater than or equal to 0.8.
<sup>b</sup>The number of instances in which the similarity score is less than 0.5.

## Conclusions

Here, we studied the effects of related fingerprints on analysis of molecular similarity by defining related fingerprints to be those that do not contribute to the shape of the eigenvalue distribution of the original fingerprint matrix. Using a dataset of human metabolites, we found that commonly used 2D structure fingerprint schemes included many related fingerprints, which led to qualitative differences in the molecular similarity analysis. Our results emphasize the potential pitfall of having highly related fingerprints for SAR analysis and suggest that an increase in the number of structural fingerprints may not always enhance the performance of molecular similarity analysis. Because our eigenvalue-based entropy approach is an unsupervised method to select related 2D fingerprints, it can be integrated seamlessly to exiting similarity-based VS pipelines which use 2D fingerprints as feature vectors.

## Acknowledgments

This work was supported by the King Abdullah University of Science and Technology (KAUST) Office of Sponsored Research (OSR) under Awards No. BAS/1/1624-01, URF/1/3412-01, URF/1/3450-01, FCC/1/1976-18, FCC/1/1976-23, FCC/1/1976-25, FCC/1/1976-26, and FCS/1/4102-02.

